# Nucleobase-Specific Recognition of DNA by Protamine

**DOI:** 10.64898/2026.11.07.765086

**Authors:** Anshika Dhiman, Hashini Ekanayake Mudiyanselage, Huan-Xiang Zhou

## Abstract

Protamines—arginine (Arg)-rich, intrinsically disordered proteins (IDPs)—replace histones to compact DNA in the sperm nucleus. While folded domains can bind DNA site-specifically, whether IDPs exhibit DNA-sequence specificity remains open. Here, we used all-atom molecular dynamics simulations to characterize protamine-DNA binding. Eight copies of salmon protamine (33 residues with 21 Arg) rapidly wrap around a 35-bp DNA molecule. In addition to salt bridges with DNA phosphates, Arg hydrogen-bonds with multiple nucleobases on both strands or the same strand. At the major groove, protamine prefers GC over AT by a 2.6-to-1 ratio, with guanine’s O6 and N7 atoms as hydrogen-bond acceptors; the preference is especially strong at G-tracts. This selectivity reverses at the minor groove, where AT overtakes GC by a 2.1-to-1 margin. The net effect is a moderate bias for GC over AT in protamine-base interactions. Additional simulations with 20-bp DNA molecules and free-energy calculations reinforce these observations. For experimental validation, we designed DNA sequences with 0–40% GC content. Protamine-DNA mixtures form aggregates, with an increasing propensity at higher GC content and upon shuffling into G-tracts. This work demonstrates that IDPs can bind DNA in a nucleobase-specific manner, with implications for protamine-mediated transcription silencing and remodeling of paternal chromatin.

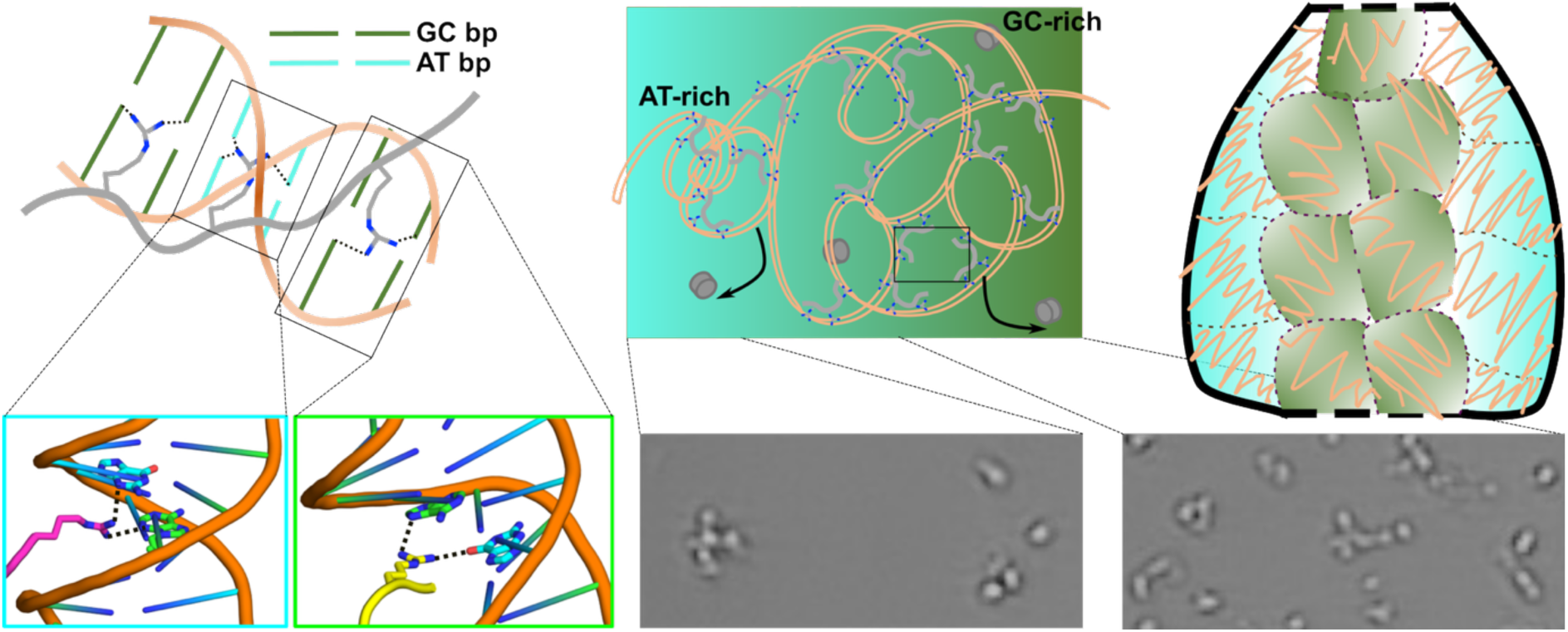

## Introduction

Protamines are small, Arg-rich, intrinsically disordered basic proteins (IDPs) [2]. In the sperm nucleus, they replace most histones and compact the chromatin approximately 10-fold more than in somatic cells [3–7]. Extreme compaction is thought to facilitate transcription silencing and protect the DNA from damage [8]. Arg residues of protamines are essential for DNA condensation as Lys substitutions are much less effective [9].

Early atomic force microscopy (AFM) of mammalian sperm chromatin and salmon protamine-DNA condensates suggested toroidal structures [10]. These observations and the notion that individual protamine molecules bind to the grooves of double-stranded (ds) DNA to neutralize the negative charges on the backbone phosphates [2,11] contributed to the development of the toroid model of sperm chromatin [3,12,13], which posits that ∼50-kb, protamine-neutralized DNA blocks spool into toroids. Recent AFM studies of protamine-bound dsDNA chains revealed a variety of structures, including stacks of loops [14], coils, rods, and doughnuts [15]. Under bulk conditions, salmon protamine condenses dsDNA into solid-like aggregates [1,16] but single-stranded (ss) DNA into liquid droplets [16,17]. Our single-molecule force spectroscopy recently demonstrated that protamine can condense dsDNA into hyperstable “tangles,” which are tightly knit bundles of loops [1]. Under mechanical tension, the forces stabilizing DNA tangles are so strong that the two strands of dsDNA separate while tangles remain intact. Coarse-grained molecular dynamics (MD) simulations suggested electrostatic interactions as the main driver for DNA condensation, with protamine-to-DNA charge ratio as the dominant determinant [18,19]. Our all-atom MD simulations showed that Arg side chains of protamine form not only salt bridges with dsDNA backbone phosphates but also hydrogen bonds with nucleobases [1]. We hypothesized that these hydrogen bonds, often in the form of wedges and clamps involving bases on opposite strands or adjacent bases on the same strand (Figure 1), respectively, may account for the hyper-stability of tangles.

**Figure 1.**
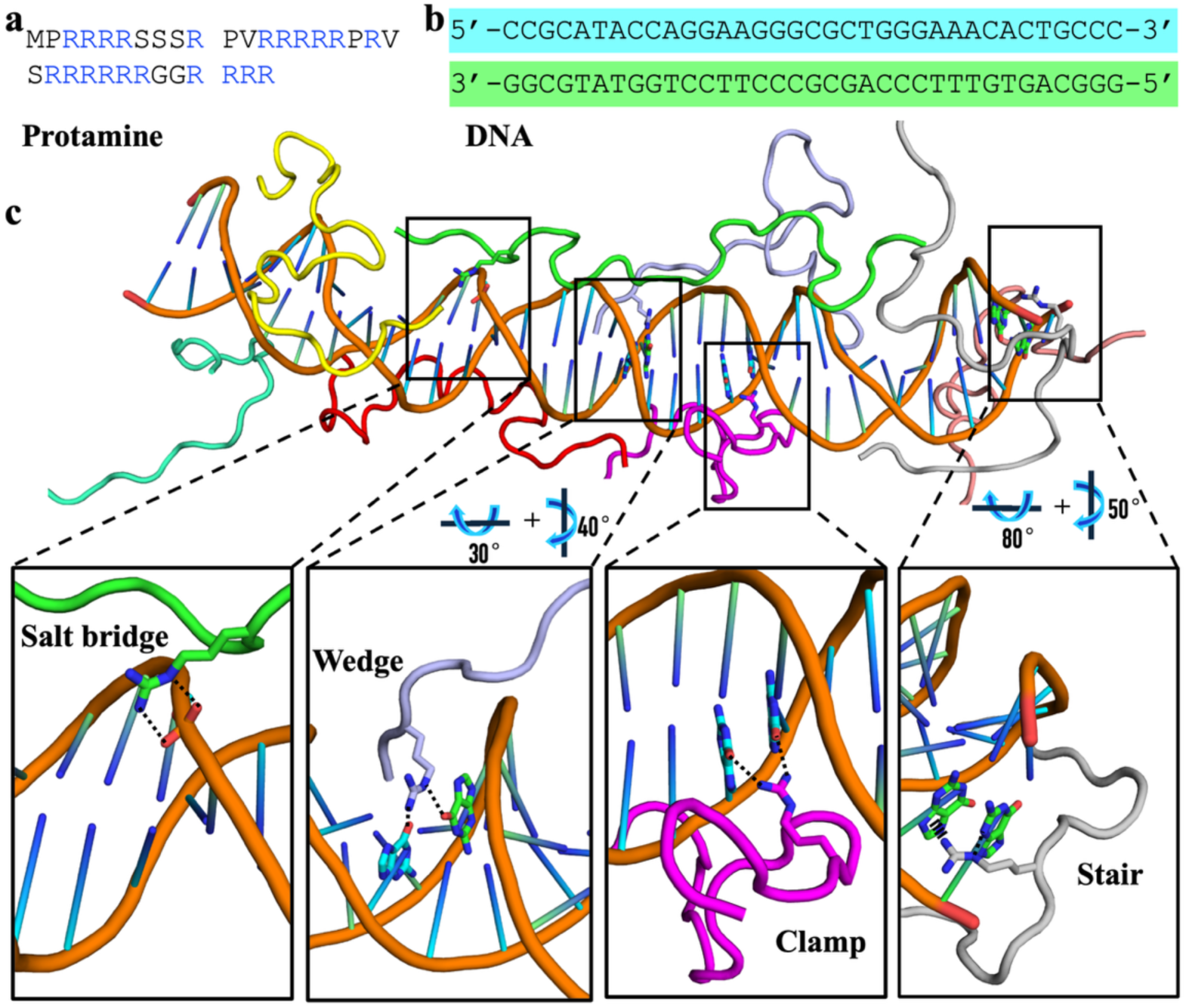
Multiple modes of side chain-nucleotide interactions. (a) Amino-acid sequence of protamine. Arg residues are highlighted in blue. (b) Nucleotide sequence of the 35-bp DNA. The two strands are colored in aqua and green, respectively. (c) A representative snapshot showing multiple modes of Arg side chain interactions with nucleotides: a salt bridge with a phosphate; a wedge comprising hydrogen bonds with bases from both DNA strands; a clamp comprising hydrogen bonds with bases in a single strand; and a stair involving both hydrogen bonding and a cation-π interaction. Eight protamine chains are shown in different colors; for the 35-bp DNA, nucleosides are displayed as sticks, with the sugar end in aqua and green to distinguish the two strands.

The code governing protein-DNA recognition has long been the object of intense scientific pursuit. Recognition can occur through direct interactions with nucleobases via hydrogen bonds, or through shape readout, in which proteins recognize sequence-dependent variations in DNA groove geometry [20]. In the major groove, the guanidinium group of Arg can form bidentate hydrogen bonds with guanine at the N7 and O6 positions, which serve as excellent hydrogen-bond acceptors [21,22]. Bidentate hydrogen bonds of Arg with two bases on the same strand or opposite strands (i.e., clamps and wedges) were observed in protein-DNA co-crystal structures [23]. In the minor groove, analysis of co-crystal structures showed that AT-rich sequences, particularly A-tracts, narrow the groove and enhance the negative electrostatic potential, facilitating preferential Arg binding [20]. However, it is unclear whether IDPs like protamines can also bind DNA in a sequence-dependent manner. Indeed, it has long been assumed that protamines bind DNA nonspecifically [2,9,11,24,25]. Free-energy simulations showed that Arg-rich peptides bind GC tracts more strongly in the major groove than in the minor groove, whereas this groove preference is reversed for AT tracts [26,27]. Using GC-only and AT-only sequences, Mandal et al. [28] further found that an Arg-rich peptide favors binding to GC tracts in the major groove but to AT tracts in the minor groove. However, the question remains open as to whether Arg-rich peptides, let alone protamines, have an overall preference for GC or AT in a mixed DNA sequence.

An inherent nucleotide-level selectivity by protamines could provide a missing mechanistic link between DNA local sequence features and global chromatin dynamics during spermiogenesis. For instance, because transcription in higher eukaryotes frequently initiates from GC-rich regions known as CpG islands [29,30], any protamine preference for GC-rich tracts could potentially interfere with transcription factor binding through steric block, thereby contributing to transcription silencing. A similar role was implicated for transition protein TP2, which is thought to facilitate the histone-to-protamine transition and exhibited a binding preference for CpG islands [31]. Moreover, such sequence-level biases may dictate genome organization. Single-cell Hi-C mapping showed that the sperm genome maintains segregated, accessible A-compartments and inaccessible B-compartments [32]; CUT&Tag mapping and ATAC-seq profiling further revealed that protamine 1 selectively loads within the accessible A-compartments before spreading globally [33]. If protamines possess an intrinsic nucleotide-level selectivity, such a bias could serve as the primary positional beacon directing this hierarchical, compartment-guided loading process, allowing the preservation of compartment segregation.

Here, we combined all-atom molecular dynamics simulations and condensate formation/dissolution experiments to investigate possible nucleobase-specific DNA binding of salmon protamine (protamine hereafter; see Figure 1a for sequence). In our simulations, eight copies of protamine bind to a 35-base pair (bp) DNA molecule with a mixed sequence (Figure 1b). The protamine molecules wrap around the DNA molecule, forming both salt bridges with backbone phosphates and hydrogen bonds with nucleobases (Figure 1c). Protamine prefers GC over AT for base interactions by a 1.3-to-1 ratio. This preference arises from an incomplete cancellation of opposite selectivity at the two grooves. At the major groove, protamine strongly favors GC over AT by a 2.6-to-1 ratio; this selectivity reverses at the minor groove, where AT overtakes GC by a 2.1-to-1 margin. Similar results are found on protamine interacting with 20-bp DNA molecules. Free energy calculations further indicate that, relative to Arg-phosphate interactions, Arg-base interactions are much stronger, making them crucial for protamine-mediated co-condensation and other DNA-related processes. To validate these results, we performed co-condensation experiments with protamine and 20-bp DNA sequences containing 0-40% GC. The mixtures form amorphous aggregates, with an increasing propensity at higher GC content and upon shuffling into G-tracts. The preference of protamine for GC, especially in G-tracts, has implications for transcription silencing and remodeling of paternal chromatin.

## Materials and Methods

### Computational: molecular dynamics simulations

All-atom simulations were performed as described previously [1], but here for a 35-bp DNA molecule at three KCl concentrations, 0, 0.15, and 1 M as well as for two 20-bp DNA molecules at 0.15 M KCl. In brief, the 35-bp DNA duplex was positioned diagonally inside a 120-Å cubic box. Eight salmon protamine molecules were then randomly placed around the duplex. The system was solvated with water and neutralized by adding 100 Cl⁻ ions as well as K^+^ and Cl⁻ ions at the desired KCl concentration. DNA, protein, water, and ions were modeled with the OL21 [34], ff14SB [35], OPC [36] force fields and the Joung and Cheatham ion parameters [37], respectively. Each system contained ∼220,000 atoms. As reference, a DNA-only system was also prepared and simulated. Similarly, systems with a 20-bp DNA molecule were prepared with five protamine molecules in a 100-Å cubic box.

The simulation of each system was initiated by 5000 steps of energy minimization (steepest descent and conjugate gradient) in AMBER22 [38], followed by a 250-ps NVT equilibration with a 1-fs timestep, during which the temperature was gradually increased from 0 K to 300 K over the first 100 ps and maintained at 300 K for the remaining duration. All heavy atoms of DNA and protamine were restrained with a 5 kcal mol⁻¹ Å⁻² force constant during the initial equilibration. An unrestrained equilibration was then performed for 1 ns under constant NPT (300 K and 1 bar) with a 2-fs time step. Production simulations were last run in four replicates for 1000 ns each under constant NPT with a 2-fs time step. All simulations were run with GPU acceleration using pmemd.cuda [39]. A cutoff of 12 Å was applied for non-bonded interactions, with long-range electrostatic interactions treated using the particle-mesh Ewald method [40]. Bond constraints involving hydrogen atoms were maintained using the SHAKE algorithm [41]. Temperature and pressure were regulated using the Langevin thermostat (damping constant 1 ps⁻¹) and Berendsen barostat [42], respectively. Snapshots were saved every 100 ps.

### Umbrella sampling

Potentials of mean force for Arg-DNA interactions were determined using a model system comprising a 7-bp DNA molecule and a single Arg side chain. The latter was built by removing the backbone atoms and capping the Cα atom with two hydrogen atoms. The resulting fragment was geometry-optimized at the HF/6-31G* level using Gaussian 16 [43]. Partial atomic charges and General AMBER Force Field 2 (GAFF2) parameters [44] were assigned using the antechamber and parmchk2 commands in Amber tools [45]. For each interaction mode, the initial structure was extracted from a snapshot in the 35-bp DNA plus protamine simulations. The model system was subjected to energy minimization and equilibration for 10 ns. Steered molecular dynamics simulations were performed in NAMD [46] to generate initial configurations for umbrella sampling. The DNA heavy atoms were restrained with a 5 kcal mol⁻¹ Å⁻² force constant while the Arg side chain was pulled away from its binding site to a 16-Å separation. Umbrella-sampling simulations were carried out in AMBER22 with a 0.25-Å interval between successive windows. These simulations lasted 10 ns, with a harmonic restraint of 250 kJ mol⁻¹ nm⁻² applied to the separation between the Arg guanidinium group and the DNA base or phosphate atoms. Meanwhile, Cartesian coordinates in the lateral directions were restrained with a 500-kJ mol⁻¹ nm⁻² force constant using PLUMED [47] to prevent lateral movement of Arg; heavy atoms were restrained with a 0.5-kcal mol⁻¹ Å⁻² to prevent DNA movement. Free energy profiles were obtained using WHAM [48].

### MD data analysis

Analyses were performed on the last 500 ns of each simulation. Protamine-DNA interactions were identified and analyzed using CPPTRAJ [49] and in-house Python/Tcl scripts, with a 3.5-Å cutoff between amino-acid side-chain heavy atoms and nucleobase polar atoms or phosphate P, OP1, and OP2 atoms. For convenience, these interactions are referred to as hydrogen bonds (bases) and salt bridges (phosphates). A wedge was assigned when one side chain hydrogen-bonds with both DNA strands, whereas a clamp was assigned when only one strand is involved. A stair was assigned when a side chain forms hydrogen bonds with one base and forms a cation-π interaction with another base. The latter was defined when the distances from Arg nitrogens to base heavy atoms were each ≤ 7 Å and differed by less than 1.5 Å, and identified using a script written by D. K. Weber (https://github.com/weberdak/flexible-interaction-tool/blob/master/flirt.tcl). When classifying bases into three modes, each base was assigned into only one mode, with the priority order wedge > clamp > stair. For example, a base that is in a wedge with one Arg was not also assigned to the clamp mode even when it is also in a clamp with another Arg. We also aggregated Arg-interactions into GC and AT subsets; their distributions were calculated using the gaussian_kde function from the SciPy library in Python.

For groove-specific analysis, major-groove atoms were N7 and O6 of guanine, N7 of adenine, and O4 of thymine; minor-groove atoms were N3 of guanine and adenine and O2 of thymine and cytosine. Radial distribution functions (RDFs) were calculated in CPPTRAJ using the radial command with a bin size of 0.05 Å. Major and minor groove widths were calculated using the 3DNA method implemented in the nastruct module of CPPTRAJ. A bp was considered partially open when any hydrogen bond therein was broken (i.e., donor-acceptor distance > 3.5 Å or donor-hydrogen-acceptor angle < 135°).

### Experimental: materials sample preparation

Salmon protamine (catalog no. P4020) and KCl (catalog no. P3911) were from Sigma (St. Louis, MO); imidazole (catalog no. AC122025000) was from Thermo Fisher. DNA constructs (see Table S1 for sequences) were from Integrated DNA Technologies (Skokie, IL).

The buffer in all co-condensation experiments was 10 mM imidazole (pH 7) plus 0.15 M KCl unless otherwise stated. Mixtures were prepared by sequentially adding DNA and protamine to the buffer and stirred well by pipetting. After mixing, samples were immediately measured.

### Brightfield microscopy

A 2-μL aliquot was placed on a glass slide and observed under an Olympus BX53 microscope with a 40× objective. Images were exported in .tiff format and processed in ImageJ.

### UV absorbance

UV absorbance was measured using a NanoDrop 2000c spectrophotometer, all in four replicates with a 2-μL sample volume each. The amount of co-condensates was monitored by the absorbance at 350 nm, which reflects light scattering by condensate particles. To measure the dilute-phase DNA concentration, samples were centrifuged at 12000 rpm for 15 min, and the supernatant was collected for absorbance measurement at 260 nm. This absorbance was converted to DNA concentration from a standard curve obtained using known DNA concentrations.

The threshold DNA concentration for co-condensation was determined using a dilution series at a fixed DNA-to-protamine molar ratio of 1:2. After a linear fit of the 350-nm absorbance, the threshold was taken as the DNA concentration where the absorbance was extrapolated to a zero value. The condensate-dissolution salt concentration was determined by measuring the 350-nm absorbance over a range of KCl concentration. The salt dependence of the absorbance was fit to the Hill equation. The dissolution salt was taken was the KCl concentration when the fit first entered background levels.

## Results

To uncover the molecular basis of protamine-DNA interactions and determine whether protamine-mediated condensation depends on DNA sequence, we performed all-atom MD simulations and co-condensation experiments.

### Arg side chains form hydrogen bonds with multiple bases

Our simulation system consists of eight copies of protamine and a 35-bp DNA molecule. Protamine is a 33-residue IDP, with 21 Arg residues (Figure 1a). The DNA sequence (derived from λ-DNA) contains 23 GC bp (66%) and 12 AT bp (Figure 1b). Simulation results at 0 M KCl were reported briefly in a previous study [1]; here we performed simulations at 0 M as well as 0.15 and 1 M KCl, each in four replicates and for 1 μs. Our main focus will be the results at 0.15 M KCl; results at 0 and 1 M KCl will be used to highlight salt effects. Within 100 ns, initially dispersed protamine molecules coalesce around the DNA. The early contacts are made primarily with the backbone phosphates, followed by interactions with the bases. We classify base interactions into three modes: wedge, clamp [1], and stair [50] (Figure 1c). They are respectively formed by an Arg side chain hydrogen-bonding to bases on both strands, to one or more bases on the same strand, and to one base while forming a cation-π interaction with another base.

We quantified protamine-DNA interactions at the residue and nucleotide levels (Figure 2). For amino-acid residues, only side chains were considered; for nucleotides, we further broke them into phosphates and bases. On average, 64.1 and 18.1 side chains (24 and 7% of all side chains or 29 and 10% among only Arg side chains) interact with phosphates and bases, respectively. Conversely, 47.0 phosphates and 31.8 bases (69 and 45% of these groups) interact with protamine side chains. These results indicate that, while side chain-phosphate interactions (in particular, salt bridges) are prevalent, side chain-base interactions (in particular, hydrogen bonds) are also substantial for the Arg-rich IDP protamine. Moreover, when side chain-base interactions occur, each side chain engages with 31.8/18.1 = 1.8 bases. In contrast, for side chain-phosphate interactions, each phosphate engages with 64.1/47.0 = 1.4 side chains. Arg accounts for 92% of all side chains that interact with bases, but only 77% of all side chains that interact with phosphates. From here on, we will concentrate on side chain-base interactions and, for simplicity, will refer to all side chains that engage in base interactions as Arg.

**Figure 2.**
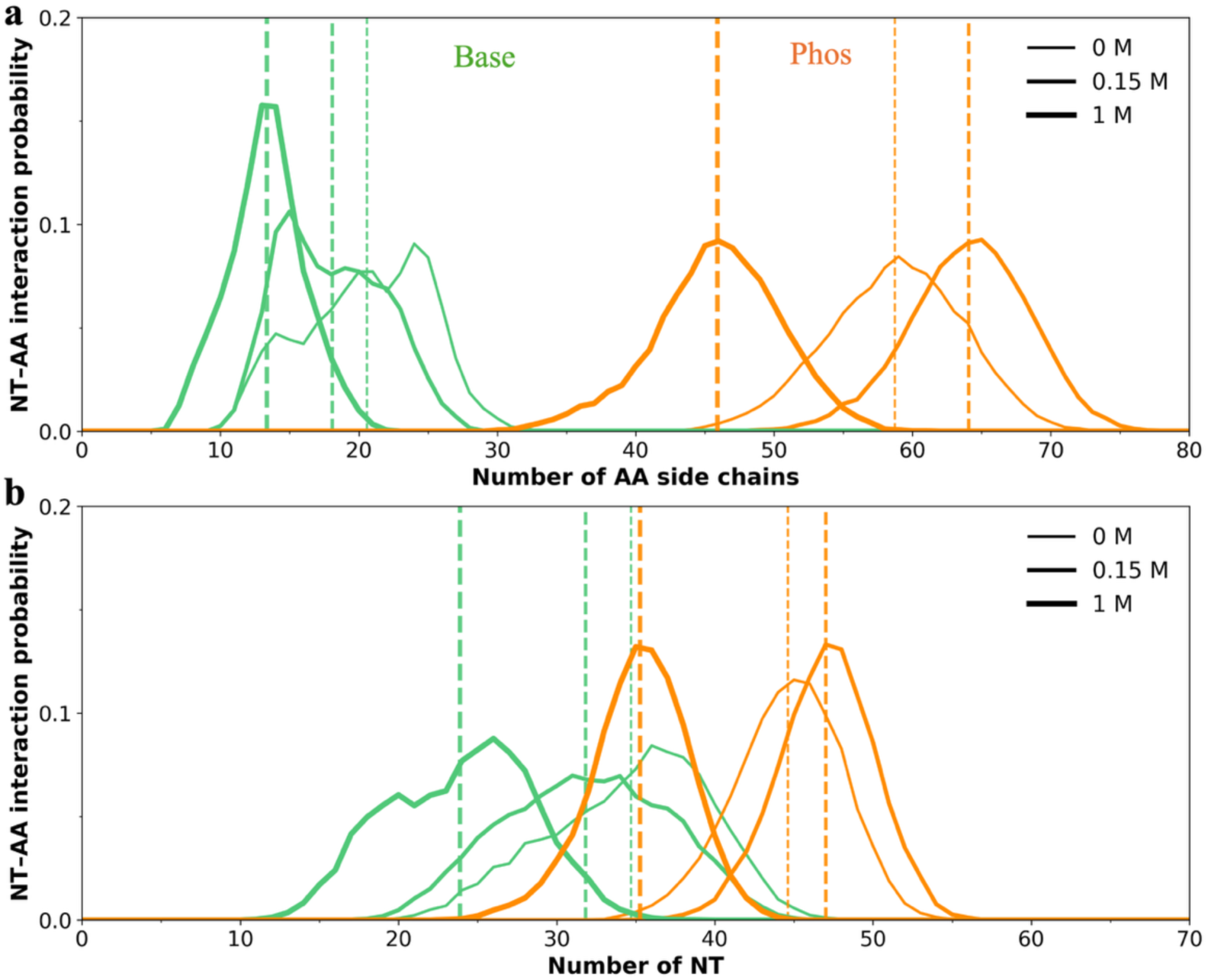
Probability distributions of protamine-DNA interactions at three salt concentrations. (a) Probability distributions of the number of protamine amino-acid side chains (AA) that interact with nucleobases or backbone phosphates. (b) Probability distributions of the number of nucleotides (NT) that interact with protamine molecules. Interactions of backbone phosphates (“salt bridges”) and bases (hydrogen bonds and cation-π) are separately enumerated. Results for three KCl concentrations are displayed as curves with increasing thickness; corresponding vertical lines indicate mean values of the distributions. Results at 0 M KCl were reported previously [1]; we reran simulations at 0 M KCl here and the results do not show significant differences: e.g., the number of base-interacting side chains was 17.3 ± 2.4 previously and is 20.6 ± 3.9 now (mean ± SD among 4 replicate simulations).

Of the 31.8 bases interacting with side chains, 23.5 are in the wedge mode and 8.3 are in the clamp mode; an additional 0.21 bases form cation-π interactions with Arg in a stair mode (Figure S1). Wedge is thus the dominant mode of Arg-base interactions, where each Arg side chain simultaneously forms hydrogen bonds with bases on both strands of the DNA molecule (Figure 1c). On the other hand, bases that only form cation-π interactions are rare in the dsDNA molecule and will not be considered further. When the KCl concentration is increased from 0 to 1 M, the counts of side chains and nucleotides forming base interactions decrease progressively, due to the disruption of hydrogen bonding by salt ions (Figure 2). Accordingly, the number of wedge-forming nucleotides declines with increasing KCl (Figure S1).

Curiously, while the counts of side chains and nucleotides forming phosphate interactions are higher at 0.15 M than at 1 M KCl, consistent with the screening of Arg-phosphate salt bridges by the salt, the counts at 0.15 M KCl are also higher than those at 0 M KCl. The maximal count of side chain-phosphate interactions at 0.15 M KCl can be attributed to the preferential localization of K^+^ ions within the major groove, due to the strong negative electrostatic potential of the DNA at that site instead of around the phosphates [51,52]. As a result, some of the Arg side chains that occupy the major groove at 0 M KCl would be displaced by K^+^ ions at 0.15 M KCl, toward nearby phosphates. The radial distribution functions of K^+^ ions provide support for this explanation (Figure S2a). The RDFs around polar atoms lining the minor groove have a higher peak at 1 M KCl than at 0.15 M KCl, in line with the expectation that the peak grows with KCl concentration. In contrast, the RDFs at the major groove run counter to this trend, with a higher peak at 0.15 M KCl than at 1 M KCl, consistent with the preferential localization of K^+^ ions within the major groove at 0.15 M KCl. Correspondingly, the RDF of Arg side chains around phosphates has the highest peak at 0.15 M KCl (Figure S2b). Cl^−^ ions are repelled from both grooves (Figure S2c).

### Arg-base interactions exhibit moderate overall preference for GC over AT

To examine possible DNA sequence specificity in protamine binding, we calculated the Arg-interaction probabilities of individual nucleobases (Figure 3). The average interaction probabilities of the four types of bases follow the order: guanine (0.67) > adenine (0.40) > thymine (0.35) > cytosine (0.32). The higher probabilities of the purines relative to the pyrimidines are expected due to their double-rings and hence greater chance of interactions with Arg, but the value for guanine is exceptionally high; guanine is unique in having two atoms, O6 and N7, on one side (i.e., the major-groove side) as hydrogen-bond acceptors to Arg (Figure 4a) [21]. Overall, GC nucleobases are favored over AT nucleobases for Arg interactions by a 1.3-to-1 ratio.

**Figure 3.**
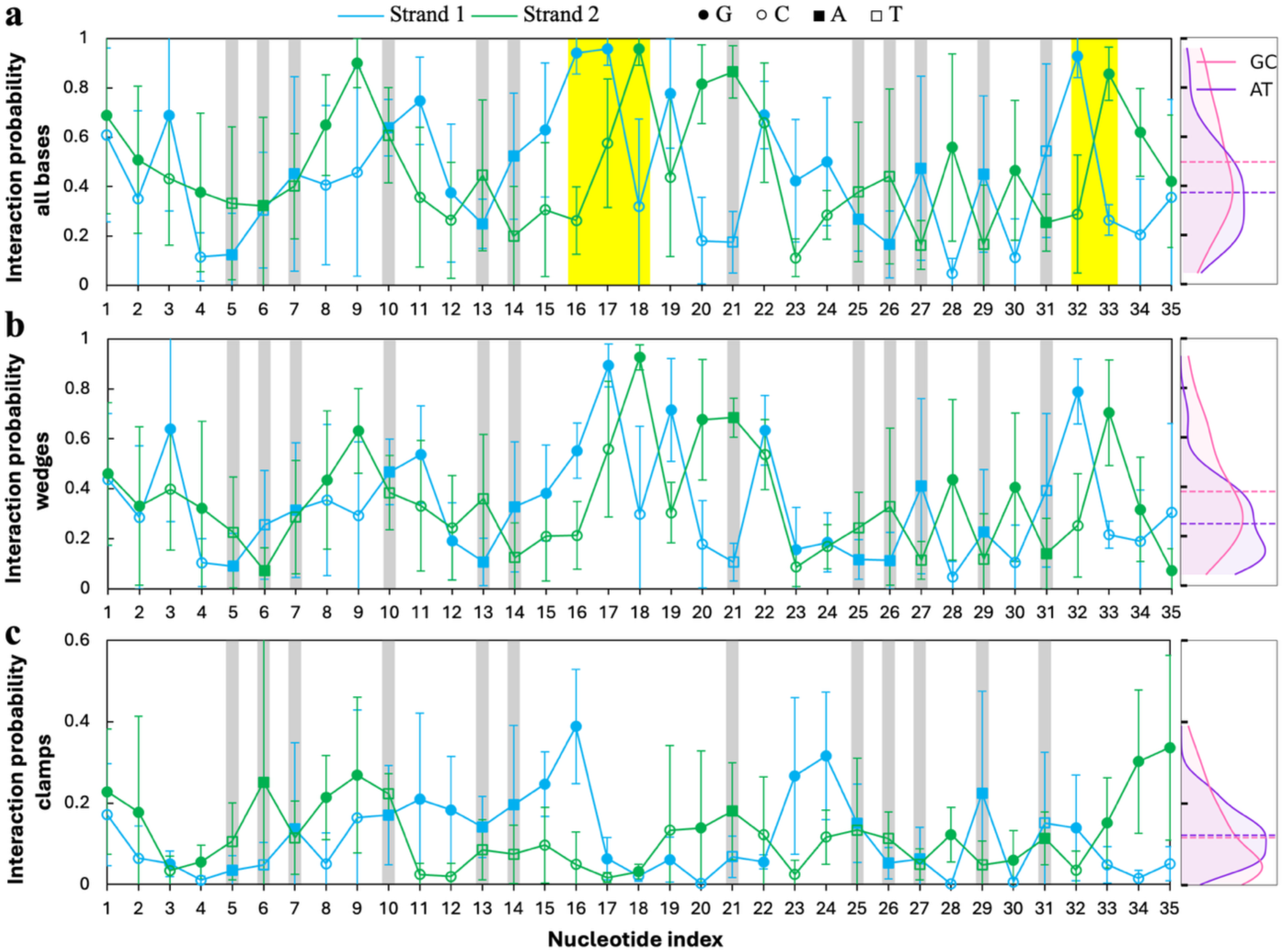
DNA sequence-specific binding of protamine. (a) Arg-interaction probabilities of bases along the two strands of the DNA; all bases included. Positions along both strands are specified by the nucleotide index along Strand 1. Results for the two strands are displayed as connected symbols in aqua and green, respectively; the symbols are filled and open circles and filled and open squares for guanine, cytosine, adenine, and thymine, respectively. Additionally, AT bp are indicated by gray shading. Yellow shading indicates upper outlying values in and around two GGG-tracts. (b) Same as (a), but for bases in the wedge mode. (c) Same as (a) but for bases in the clamp mode. Error bars represent SDs among four replicate simulations. Side histograms show the distributions of interaction probabilities for guanine and cytosine as a subset (GC; pink) and adenine and thymine as a subset (AT; purple). Horizontal lines show mean values in each subset.

**Figure 4.**
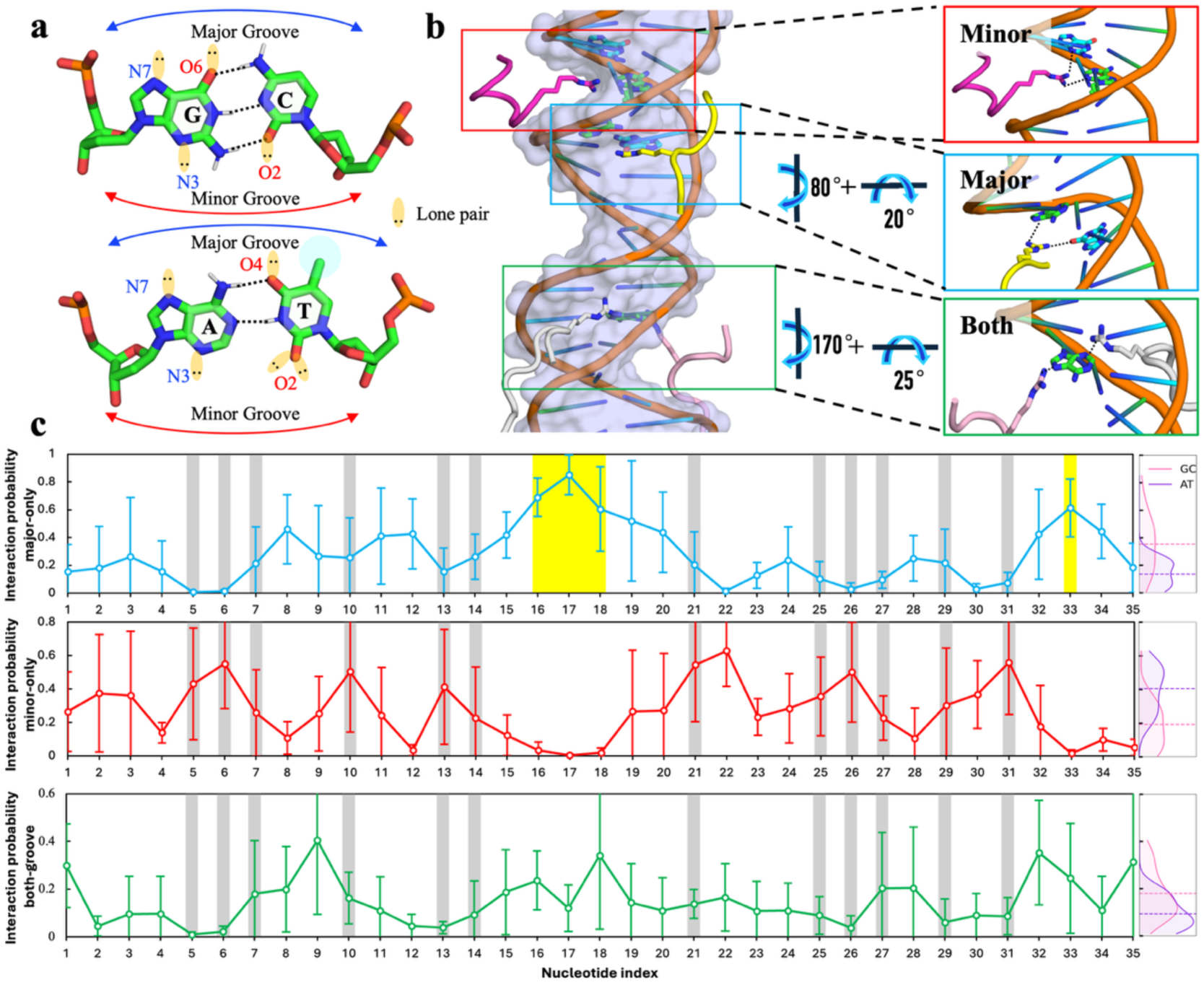
Contrasting bp preferences of protamine at the major and minor grooves. (a) Chemical structures of the GC and AT bp, highlighting the polar atoms that can serve as hydrogen-bond acceptors to Arg at the major and minor grooves. (b) A snapshot showing bp forming hydrogen bonds with Arg at the minor groove only, the major groove only, and both grooves. (c) Arg-interaction probabilities of bp along the DNA sequence for the major-only (top), minor-only (middle), and both-groove (bottom) groups. Error bars represent SDs among four replicate simulations. Side histograms show the distributions of interaction probabilities in each group when the bp are further divided into GC (pink) and AT (purple) subsets. Horizontal lines show mean values in each subset.

Figure 3a displays the interaction probabilities of nucleobases along the DNA sequence. For each of the 23 GC bp, the interaction probability of guanine is higher than that of cytosine; in contrast, for only slightly more than half (7 out of 12) of the AT bp, the interaction probability of adenine is higher than thymine. This contrast again highlights the unique ability of guanine as hydrogen-bond acceptors to Arg. Among GC nucleobases, interaction probabilities are uneven. Using mean + 1.5 SD (calculated among all GC bases) as the threshold, two stretches, at positions 16-18 and 32-33 along Strand 1, stand out as upper outliers (indicated by yellow shading in Figure 3a). These stretches overlap with two of the three GGG-tracts in the DNA sequence, indicating cooperative effects in G-tracts. No similar outliers exist among AT nucleotides, indicating more even interaction probabilities.

We separate Arg-interacting nucleotides according to modes, i.e., wedge and clamp, and display interaction probabilities of nucleobases in these modes in Figure 3b, c. Among wedge-mode nucleobases, GC is favored over AT by a 1.5-to-1 margin, higher than the total population. In comparison, clamp-mode nucleobases do not show a bias between GC and AT. The overall preference of GC over AT is preserved as the KCl concentration changes from 0 to 1 M (Figure S3). At 0 M KCl, the interaction probabilities at the third GGG-tract (positions 22-24) are also upper outliers.

### Protamine preferentially binds to GC in the major groove but AT in the minor groove

Since each Arg side chain can interact with a bp from only one of the two grooves of the DNA (Figure 4a), we break Arg-interacting bp into groups where interactions occur at the major groove only, the minor groove only, and at both grooves simultaneously (Figure 4b). All three groups are well populated, indicating no groove preference *per se*. However, within each group, Arg-interaction probabilities show substantial biases between GC and AT (Figure 4c). In the major-only group, average interaction probabilities of GC and AT are 0.355 and 0.136, respectively, resulting in a 2.6-to-1 preference for GC.

As already noted, the preference of Arg for guanine at the major groove can be attributed to the fact that two of its atoms, O6 and N7, can serve as hydrogen-bond acceptors. RDFs of Arg around these atoms indeed show very high peaks (Figure S4a). Cytosine does not have any hydrogen-bond acceptor at the major groove; the only polar group there, an amine, can only serve as a donor. Still, the RDF of Arg around this atom has a moderate peak (Figure S4b). This peak is due to a “bystander” effect: as cytosine amine nitrogen (N4) is hydrogen-bonded to guanine O6 in a bp, an Arg side chain that hydrogen-bonds with guanine O6 tends to have close contact with cytosine N4 (Figure S4c). The RDFs of Arg around adenine N7 and thymine O4 have moderate peaks (Figure S4d, e). The tempered peak for thymine O4 is due to the neighboring methyl acting as a steric blocker [53]. The RDF for adenine amino nitrogen (N6) also has a moderate peak, thanks to its proximity to N7 on the same base (Figure S4f) and to O4 in the partner thymine base (Figure S4g). Conversely, this proximity, for steric reasons, may interfere with hydrogen bonding of Arg with the N7 and O4 atoms. In short, the preference of Arg for GC over AT at the major groove stems from the guanine O6 and N7 atoms both as hydrogen-bond acceptors as well as from the thymine methyl and adenine amino groups as steric blockers.

In the minor-only group, average interaction probabilities of GC and AT are 0.193 and 0.405, respectively, corresponding to a 2.1-to-1 preference for AT (Figure 4c middle row). At the minor groove, RDFs of Arg around adenine N3 and thymine O2 both have very high peaks (Figure S4d, e). In comparison, the RDF peak around cytosine O2 is high (Figure S4b) and that around guanine N3 is moderate (Figure S4a). The tempered peaks may be due to the proximity to (and hence steric hindrance by) the guanine 2-amino group [54] (Figure S4h). Additionally, groove shape may contribute to the AT preference [20]. AT can narrow the minor groove and generate a stronger negative electrostatic potential than GC. Compared to naked DNA, our simulations show that protamine binding produces narrowing of both grooves (Figure S5): while the major groove narrowing is modest and has no overt nucleotide dependence, the minor groove narrowing is prominent, especially at AT bp. The bulky amino group is thought to underlie guanine’s resistance to minor groove narrowing [54,55].

The stronger preference for GC over AT at the major groove than the opposite preference at the minor groove produces the moderate net bias toward GC reported in the preceding subsection. For nucleobases that interact with Arg at both grooves, the average interaction probabilities of GC and AT are 0.180 and 0.093, respectively, again pointing to GC as the favored target of Arg (Figure 4c bottom row). Upon protamine binding, AT but not GC bp have elevated tendencies to be partially open (Figure S6). Arg side chains can outcompete the AT base partners for hydrogen bonding, but are unable to perturb the stronger GC bp. Put differently, with GC, edge-on hydrogen bonds are stable without partial bp opening, but with AT, edge-on hydrogen bonds may be stabilized by partial bp opening.

We highlighted the cooperative effects in G-tracts (Figure 3a, yellow shading); now we can see these effects manifest at the major groove, as indicated by the outlying Arg-interaction probabilities of bp at positions 16-18 and 33 (Figure 4a top row, yellow shading). Although the bp at position 32 is not an upper outlier in the major-only group, it is so in the both-groove group (Figure 4c bottom row). A GC bp can readily switch from the both-groove group to the major-only group by breaking interactions at the minor groove (see next paragraph). A G-tract increases the residence time of an Arg side chain at the major groove, as the side chain shifts its grip from one guanine to the next as if traversing monkey bars (Movie S1). (The monkey-bar analogy has been used previously for traversing different DNA segments [56].)

Over the KCl concentration range from 0 to 1 M, the total interaction counts decrease progressively, due to salt screening (Figure S7). From 0 to 0.15 M KCl, the mean interaction probabilities hold steady in the minor-only group, decrease in the both-groove group, but actually increase in the major-only group. The latter increase can be attributed to the GC bp that form Arg interactions at both grooves at 0 M KCl but break such interactions at the minor groove at 0.15 M KCl, leading to a conversion from the both-groove group to the major-only group. Arg-GC interactions at the major groove have greater ability to withstand salt screening than at the minor groove.

### Simulation results with 20-bp molecules confirm nucleobase- and groove-specific recognition of Arg

To confirm the generality of the foregoing findings, we performed additional simulations where a 20-bp DNA molecule is surrounded by five copies of protamine. This DNA molecule has a 40% GC content (Figure S8a) and is a part of the series for which we have performed co-condensation experiments with protamine (Table S1). The results are largely similar to those for the 35-bp DNA. For example, when protamine-DNA interactions form, each side chain engages with 1.6 bases but each phosphate engages with 1.2 side chains (Figure S8b, c), verifying the ability of the Arg side chain to hydrogen bond with multiple bases, with wedge as the dominant mode (Figure S8c inset). Likewise, GC in the 20-bp DNA exhibits a 1.3-fold higher mean interaction probability than AT (Figure S9).

Regarding groove specificity, in the major-only group for the 20-bp DNA, the mean interaction probability of GC is higher than the counterpart of AT by a 1.5-to-1 ratio (Figure S10 top row), whereas in the minor-only group, the trend is reversed, with AT favored by a 1.4-to-1 margin (Figure S10 middle row). Both of these margins are lower than the counterparts for the 35-bp DNA. In the major groove, the muted GC preference comes directly from a much reduced interaction probability of GC (from 0.355 to 0.155) on going from the 35-bp DNA to the 20-bp DNA, which in turn can be attributed to the lack of any G-tract in the 20-bp sequence. In the minor groove, the tempered AT preference mostly arises from an elevated interaction probability of GC (from 0.193 to 0.282) instead of a reduced interaction probability of AT. In particular, three isolated GC bp, at positions 7, 9, and 17 of Strand 1, benefit from the high interaction probabilities of neighboring AT bp. Therefore, high interaction probabilities can spill over into a neighboring bp. For bases that interact with Arg at both grooves, the average interaction probabilities of GC and AT are 0.218 and 0.087, respectively (Figure S10 bottom row); these values are similar to their counterparts in the 35-bp DNA and confirm GC as the favored target of Arg.

We further inquired the reason for the lower GC preference at the major groove of the 20-bp DNA relative to the counterpart of the 35-bp DNA. Inspection of their sequences revealed that the 20-bp DNA lack the G-tracts that are important for elevating the Arg interaction probabilities at the major groove of the 35-bp DNA. To test this role of G-tracts, we shuffled the eight guanines (present as four singles and two doubles) in the 20-bp DNA into two GGG-tracts and two singles (Figure S11a; Table S1). To distinguish the two DNA sequences, we refer to the one before shuffling as 40% and the after shuffling as 40%t (“t” for G-tract). The protamine interaction characteristics of 40%t are largely similar to those of 40%, including the engagement of 1.8 bases per bound side chain (Figure S11b, c) and an overall 1.4-to-1 preference of GC over AT (Figure S12). However, the preference for GC at the major groove of 40%t is elevated to 2.7-to-1, and meanwhile the preference for AT at the minor groove is elevated to 1.8-to-1 (Figure S13). Moreover, just as we expect, the elevation in the preference for GC at the major groove is largely due to the two G-tracts. This assertion is evidenced by the outlying values of Arg-interaction probabilities at positions 4-5 and 13-14 along Strand 1, which are within the two G-tracts (yellow shading in Figure S12a), and by the outlying values at these positions in the major-only (yellow shading in Figure S13 top row) and both-groove groups. In short, G-tracts are effective in promoting Arg interactions.

### Arg interactions with bases are much stronger than with phosphates

To quantitatively compare the energetic contributions of Arg-DNA interactions via nucleobases vs. via backbone phosphates, we performed free-energy calculations (Figures 5 and S14). For this purpose, we used a 7-bp model for DNA and a hydrogen-capped side chain for Arg. We then performed 1D umbrella sampling by pulling the Arg side chain away from an initial wedge (Figure 5a), clamp (Figure 5c inset), or salt-bridge pose (Figure 5d inset). The resulting free-energy profiles are displayed in Figures 5b-d and S14. The wedge mode involving two guanines on opposite strands and at the major groove is the strongest, with a free energy of –4 kcal mol⁻¹ (Figure 5b), followed by the clamp mode involving two guanines on the same strand at –3 kcal mol⁻¹ (Figure 5c). The salt bridge with a phosphate is the weakest, with a free energy of –1.3 kcal mol⁻¹ (Figure 5d). Even though the number of phosphates interacting with protamine is higher than the number of nucleobases (by < 1.5-fold), wedges and clamps still make the dominant contributions to protamine-DNA interaction stability. All these modes of Arg interactions are stronger at 0 M KCl and weaker at 1 M KCl, as can be expected from salt screening.

**Figure 5.**
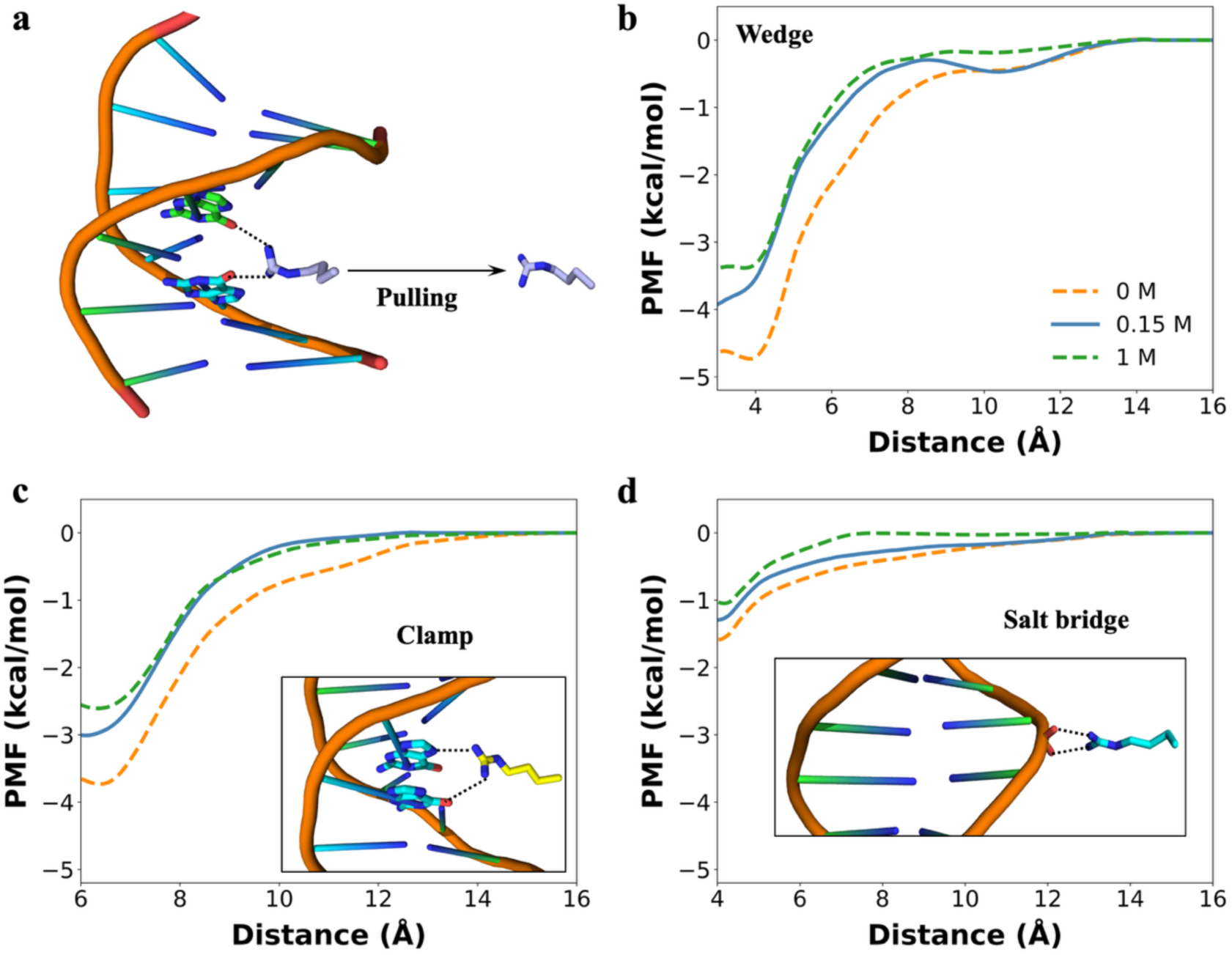
Potentials of mean force (PMFs) for Arg-DNA interactions. (a) A schematic showing an Arg side chain is pulled away from a wedge mode. (b) PMF profiles for the wedge mode, involving two guanines at the major groove, at three KCl concentrations. (c) Counterparts for the clamp mode. (d) PMF profiles for a salt bridge with a backbone phosphate. In (c, d), insets display the corresponding interactions.

We also calculated the free-energy profiles for a wedge and a clamp involving two adenines (Figure S14). In qualitative agreement with the preference of Arg for GC over AT at the major groove, the wedge and clamp involving two thymines are less stable than those involving two guanines, with free energies at –2.2 and –1.5 kcal mol⁻¹, respectively. Unfortunately, due to its narrowness, no path exists that leads to the breakup of a wedge or clamp without interference by backbone phosphates and corresponding free-energy calculations at the minor groove could not be made. Together, the conventional and free-energy MD simulations allow us to conclude that the strength and valency of protamine-DNA interactions should increase with an increasing GC content and upon shuffling into G-tracts, leading to an elevated propensity for co-condensation [57].

### Higher GC content and G-tracts promote protamine-DNA co-condensation

To validate the foregoing conclusion, we designed four 20-bp DNA sequences, one each at 0 and 25% GC and two at 40% GC (Table S1); in the latter two sequences, the eight guanines are either dispersed in Strand 1 or partially clustered into two GGG-tracts. We performed co-condensation experiments with protamine and characterized the differences in co-condensation among the four DNA sequences in several ways. As reported previously, protamine condenses dsDNA into aggregates [1,16,58]. Brightfield images show that, at a constant 20-bp DNA and protamine concentrations of 10 μM and 20 μM, respectively, the co-condensation propensity rises with higher GC content as indicated by an increasing number of aggregate particles, though the difference between 40% and 40%t are hard to discern (Figure 6a). We then quantified the total amounts of aggregates by measuring absorbance at 350 nm (A_350_; Figure 6b). The A_350_ values exhibit a steady rise as the GC content is increased and a small further rise as guanines are shuffled into G-tracts.

**Figure 6.**
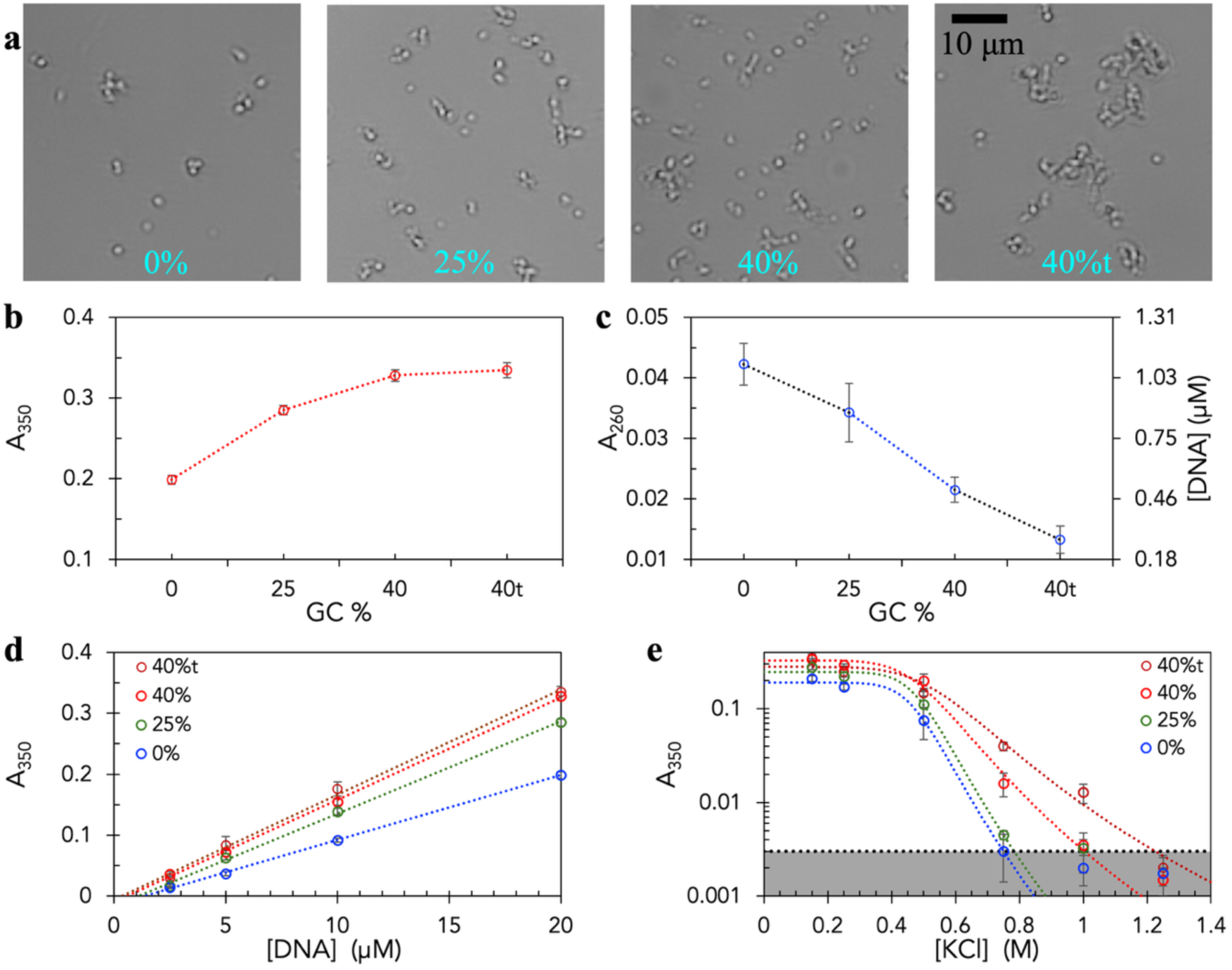
Dependence of protamine-DNA co-condensation on GC content and guanine positioning along the DNA sequence. (a) Brightfield images of DNA-protamine co-condensates, formed by mixing 10 μM 0%-, 25%-, 40%-, and 40%t-GC DNA with 20 μM protamine. (b) Amounts of co-condensates, measured by absorbance at 350 nm (A_350_), formed by mixing 20 μM DNA with 40 μM protamine. (c) Dilute-phase DNA concentrations, measured by supernatant absorbance at 260 nm (A_260_) after centrifugation. (d) Threshold DNA concentrations for co-condensation with protamine, measured by A_350_ in a dilute series where the DNA-to-protamine molar ratio is fixed at 1:2. Lines show linear fits. (e) Salt dissolution of co-condensates formed by mixing 20 µM DNA with 40 µM protamine. Curves show fits to a Hill equation; the shaded region indicates background levels. In (b-e), Error bars represent SDs among four replicate measurements. The buffer was 10 mM imidazole (pH 7) plus 0.15 M KCl (or as otherwise indicated).

In addition, we centrifuged the samples to collect the supernatant to measure the DNA levels in the dilute phase using absorbance at 260 nm (A_260_; Figure 6c). Corresponding to the increasing amounts of aggregates, the dilute-phase DNA levels, also known as saturation or threshold concentrations, show a gradual decline at higher GC content and with shuffling into G-tracts. By determining a standard curve (Figure S15), the threshold DNA concentrations are 1.1, 0.9, 0.5, and 0.3 μM, respectively, for the 0%-, 25%-, 40%-, and 40%t-GC sequences. We also obtained the threshold DNA concentrations in an alternative way, by tracking the decrease in A_350_ in a dilution series (Figure 6d). By extrapolating to A_350_ = 0, the threshold DNA concentrations are 1.3, 1.1, 0.6, and 0.3 μM for the four DNA sequences, which are comparable to those measured after centrifugation and confirm the increasing trend in co-condensation propensity.

Lastly, we used salt to dissolve protamine-DNA aggregates (Figure 6e). In line with the decreased stability of protamine-DNA interactions in the conventional and free-energy MD simulations, the amount of aggregates formed with each DNA sequence steadily decreases at increasing KCl. Moreover, the KCl concentrations at which aggregates are fully dissolved, 0.76, 0.79, 1.0, and 1.2 M, respectively, for the 0%-, 25%-, 40%-, and 40%t-GC sequences, are in complete accord with the co-condensation propensities.

## Discussion

By combining MD simulations and co-condensation experiments, we have characterized the atomic details of protamine-DNA interactions, in particular arginine side chain-nucleobase hydrogen bonding, and their roles in co-condensation propensity and condensate stability. We previously hypothesized that such hydrogen bonding, manifested as wedges and clamps, may account for the hyper-stability of tangles observed in single-molecule experiments [1]. The results presented here strongly support this hypothesis. Moreover, we show that Arg-nucleobase hydrogen bonding overall prefers GC over AT, and this preference arises from incomplete cancelation of a very strong bias for GC at the major groove and a less strong bias for AT at the minor groove. Lastly, the preference of GC over AT is promoted by the clustering of guanines into G-tracts. Together, these results have profound implications for transcription silencing in mature sperm and chromatin remodeling during spermiogenesis.

Our work counters the long-held assumption that protamine binds DNA nonspecifically [2,9,11,24,25] and shakes the foundation—the notion that the main role of protamine is to neutralize the negative charges on the backbone phosphates [2,11]—of the classical toroid model of sperm chromatin [3,12,13]. Although salt bridging of Arg with DNA backbone phosphates may be prevalent, our conventional and free-energy simulations reveal that hydrogen bonding with nucleobases is more energetic and therefore crucial for protamine-mediated DNA condensation. The crucial role of Arg-base hydrogen bonding is directly validated by the experimental data showing the rises in both co-condensation propensity and resistance of condensates to salt dissolution at higher GC content and upon shuffling into G-tracts.

Our finding that protamine favors GC over AT in DNA binding provides a simple explanation for the differential competition between protamine and GC-selective vs. AT-selective dyes for sperm DNA reported by Bizzaro et al. [59] These authors performed a competition assay between salmon protamine and a GC-selective (chromomycin A_3_) or AT-selective dye (DAPI) on decondensed mouse or human sperm nuclei where endogenous protamines were removed by high salt. Protamine recondensed the sperm nuclei and completely inhibited the binding of chromomycin A_3_, but the inhibition was only 50% for DAPI. The differential inhibitory effects are in perfect accord with the preferentially binding of protamine to GC over AT. Preferentially binding of protamine to GC-rich regions may repress transcription initiation at CpG islands, as was suggested for TP2 [31]. Moreover, as GC-tracts at the major groove are often the recognition sites of transcription factors and other regulatory proteins [60], preferential binding at these sites by protamine may block such recognition. Therefore, protamine may drive transcription silencing in sperm nuclei not only through global DNA compaction but also through preferential binding to GC-rich regulatory elements.

Recent CUT&Tag mapping revealed that mouse protamine 1 selectively loads within the accessible A-compartments before spreading globally [33]. Given that A-compartments are relatively GC-rich, protamine’s selective loading there is naturally explained by its preference for GC found here. Also, nucleosomes show a higher intrinsic occupancy in GC-rich regions [61], which dictates their selective retention at such loci in mature sperm [62]. Competition by protamine for GC-rich regions may aid in nucleosome eviction during histone-to-protamine replacement. This idea is supported by three lines of evidence. First, an *in vitro* competition assay has demonstrated that protamine can disassemble nucleosomes and displace histones [63]. Second, protamine knockout leads to elevated histone retention [64]. Third, exogenous protamine expression in somatic cells produces partial histone displacement [65].

In conclusion, we find that protamine favors GC over AT at the major groove but AT over GC at the minor groove, with a moderate overall preference for GC (Figure 7a). In genomic DNA, sequence composition dictates polymer mechanics and nucleosome retention: flexible, AT-rich regions are inherently more prone to collapse, whereas rigid, GC-rich regions are more open while retaining a fraction of nucleosomes (Figure 7b). These physical differences lead to differential condensation by protamine and ultimately underlie the preservation of sperm chromatin compartmentalization [32] (Figure 7c).

**Figure 7.**
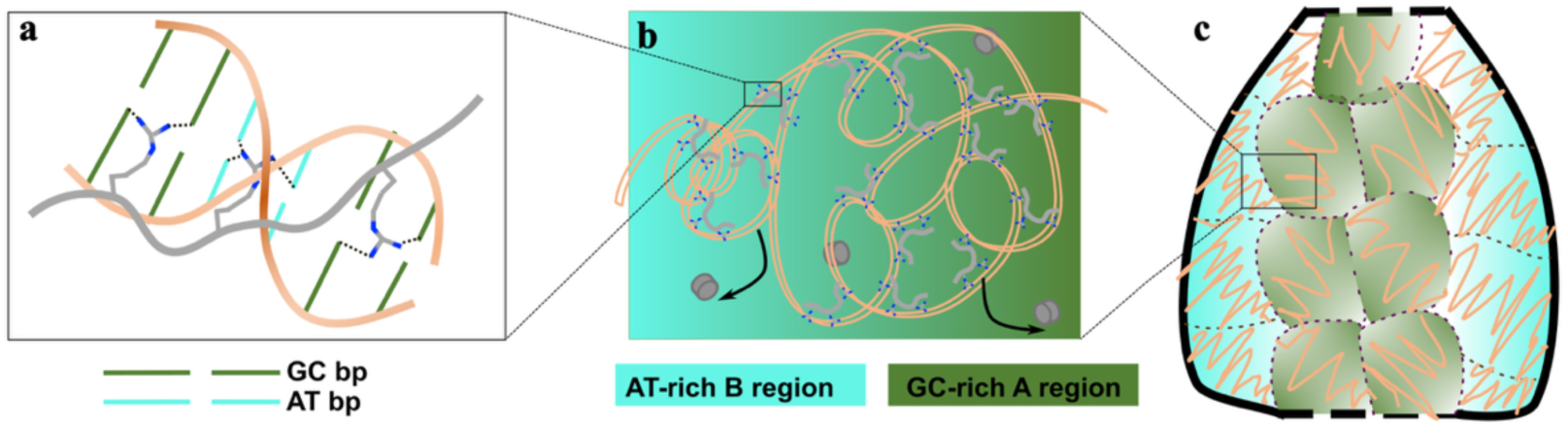
Nucleobase-specific recognition of DNA by protamine and implication for spermiogenesis. (a) Arg-nucleotide hydrogen bonding with GC at the major groove and AT at the minor groove. (b) Differential DNA chain collapse, nucleosome retention, and condensation by protamine in AT-rich B regions and GC-rich A regions. (c) Compartmentalization of sperm chromatin.

## Data Availability

Source data are deposited in GitHub (https://github.com/hzhou43/protamine-DNA).

## Supplementary Data

Supplementary Table S1, Figures S1-S15, and Movie S1.

## Author Contributions

Conceptualization: A.D., H.E.M., and H.-X.Z. Methodology: A.D. and H.E.M. Investigation: A.D., H.E.M., and H.-X.Z. Funding acquisition, project administration, supervision: H.-X.Z. Writing: A.D., H.E.M., and H.-X.Z.

