## Supplementary Table and Figures for "Nucleobase-Specific Recognition of DNA by Protamine"

*Supplementary Materials*

**Table S1. Designed 20-bp sequences.**

| Name / GC content (%) | Sequence |
|---|---|
| 0 | 5′-AAAAAAAAAAAAAAAAAAAA-3′ |
| 25 | 5′-AAA**G**AAAAA**G**AA**G**AAA**G**A**G**A-3′ |
| 40 | 5′-AAA**G**AA**G**A**G**A**GG**AAAA**G**A**GG**-3′ |
| 40t | 5′-AAA**GGG**AAA**G**AA**GGG**AA**G**AA-3′ |

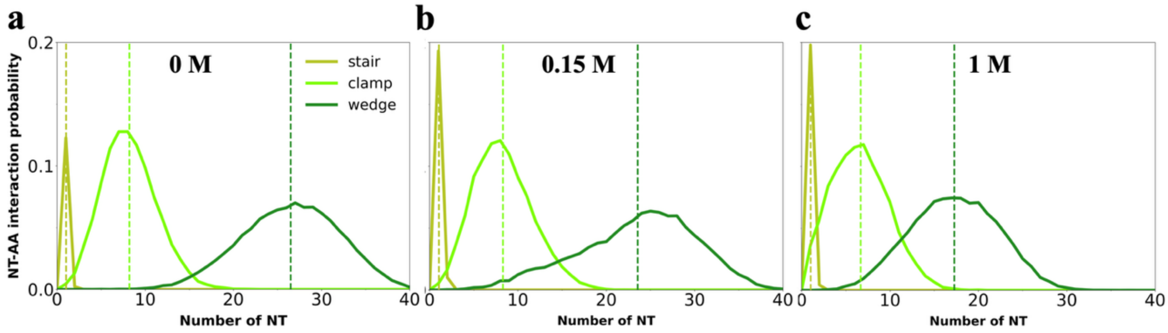


**Figure S1. Probability distributions of the number of Arg-interacting nucleotides at three salt concentrations.** (a-c) Results at 0, 0.15, and 1 M KCl. Nucleobases are separated according to their Arg-interacting modes: stair, clamp, and wedge. Dashed vertical lines indicate the mean number of Arg-interacting nucleotides in each mode.

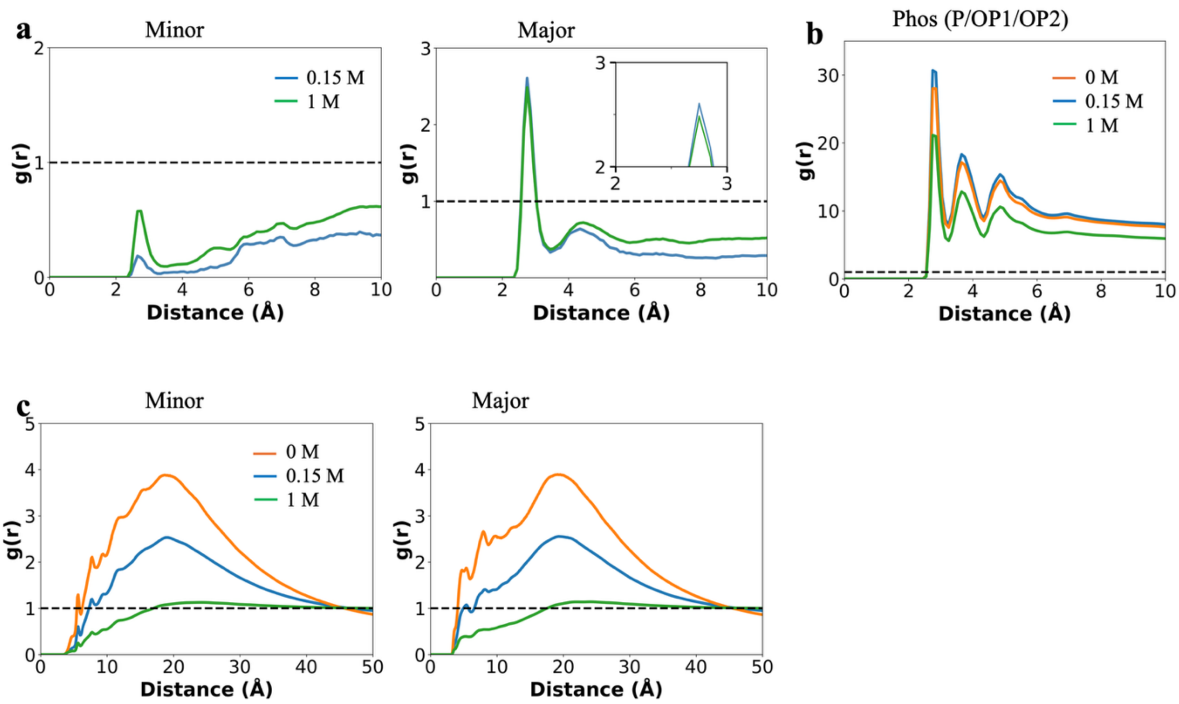


**Figure S2. Radial distribution functions [g(r)] of ions and Arg around DNA sites.** (a) g(r) for $K^+$ ions around the minor and major grooves at 0.15 and 1 M KCl. (b) g(r) for Arg side chains around backbone phosphates at 0, 0.15, and 1 M KCl. (c) g(r) for $Cl^-$ ions around the minor and major grooves at 0, 0.15, and 1 M KCl. Dashed horizontal lines are drawn at g(r) = 1.

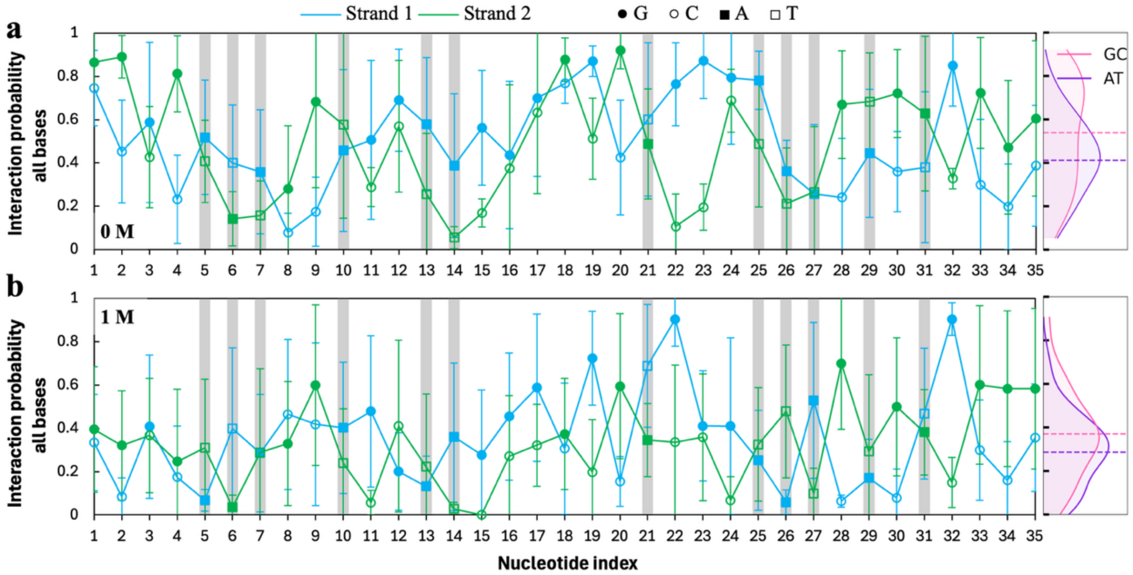


**Figure S3. DNA sequence-specific binding of protamine.** Same as Figure 3, but for KCl concentrations at (a) 0 M and (b) 1 M KCl.

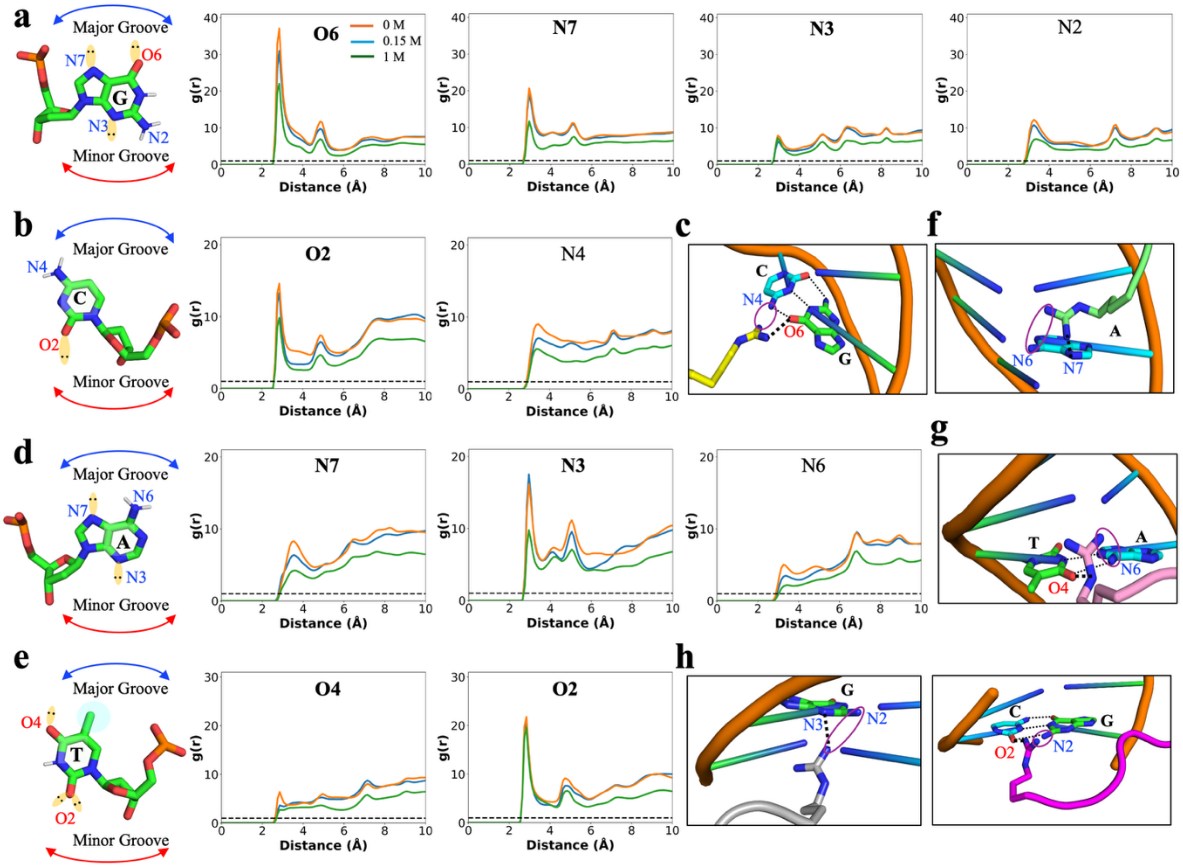

**Figure S4. Radial distribution functions [g(r)] of Arg around DNA nucleobase atoms at three salt concentrations.** (a) g(r) around guanine polar atoms. (b) g(r) around cytosine polar atoms. (c) A snapshot showing the proximity of cytosine N4 (highlighted by a magenta oval) to Arg when the latter hydrogen bonds to the bp partner guanine O6 (indicated by thick dash). (d) g(r) around adenine. (e) g(r) around thymine. (f-h) Similar to (c), but for proximity to other amino nitrogens.

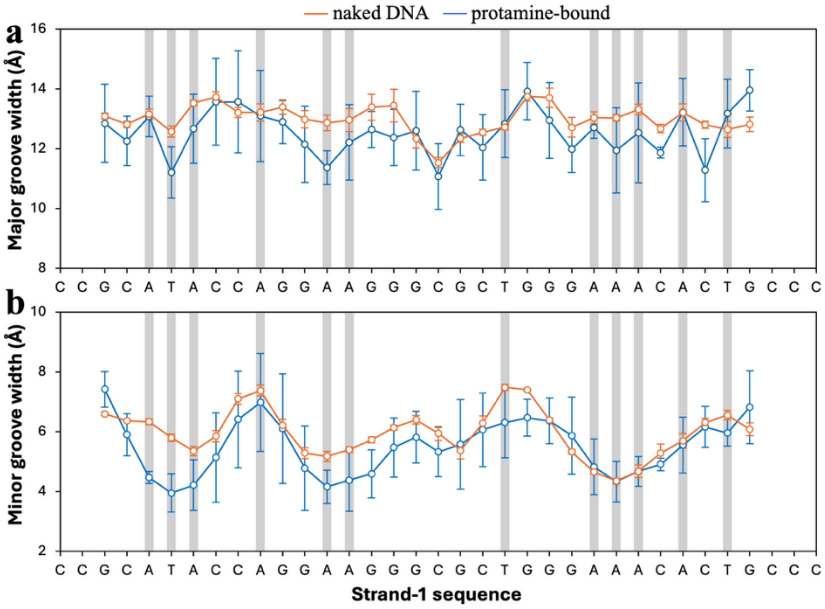


**Figure S5. Groove widths of naked DNA and after binding protamine.** (a) Major groove. (b) Minor groove. Error bars represent SDs among four replicate simulations.

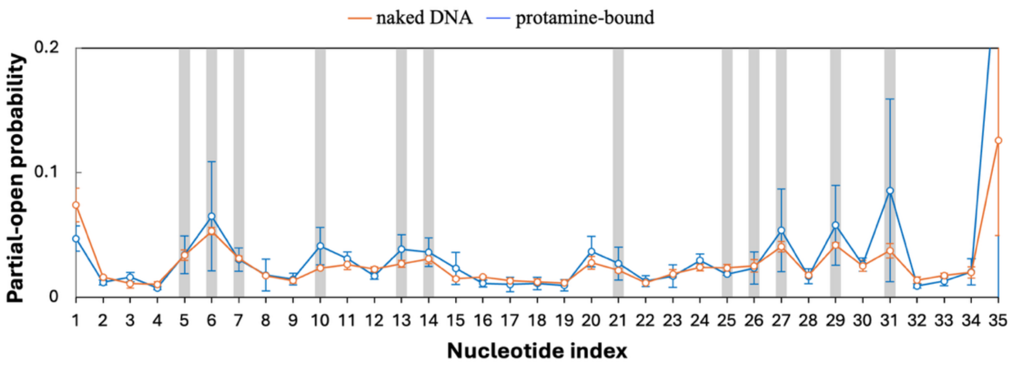


**Figure S6. Base-pair partial-opening probabilities of naked DNA and after binding protamine.** Error bars represent SDs among four replicate simulations.

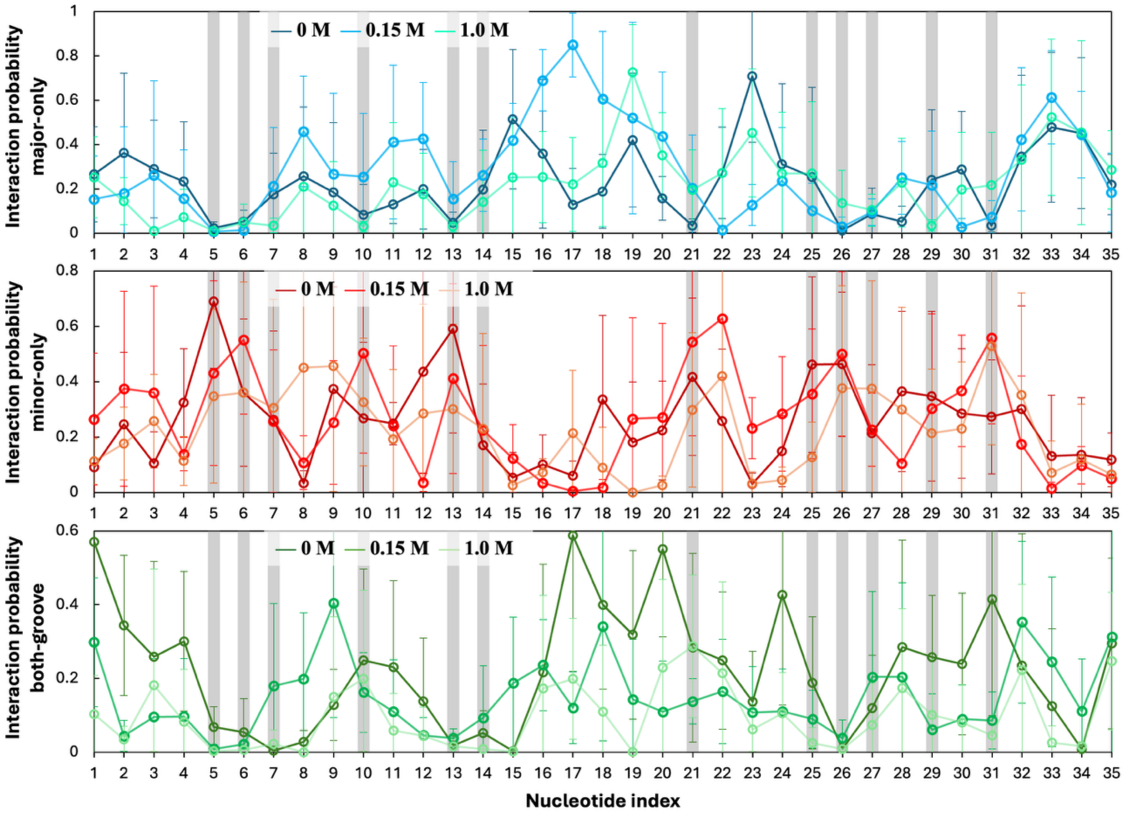


**Figure S7. Groove-specific Arg-interaction probabilities of bp along the DNA sequence.** Same as Figure 4c, but with results at 0 and 1 M KCl added.

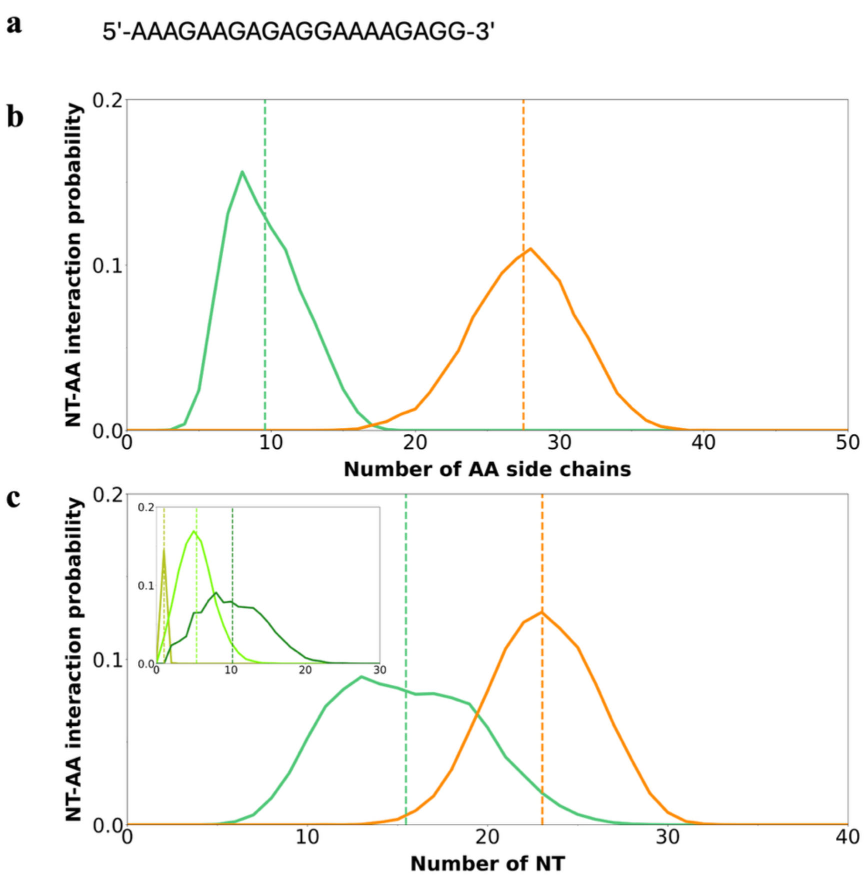


**Figure S8. Probability distributions of protamine interactions with 20-bp DNA.** (a) Sequence of the 20-bp DNA with 40% GC content. (b, c) Same as Figure 2a, b but for the 20-bp DNA and at 0.15 M KCl. The inset in (c) is the same as Figure S1b but for the 20-bp DNA.

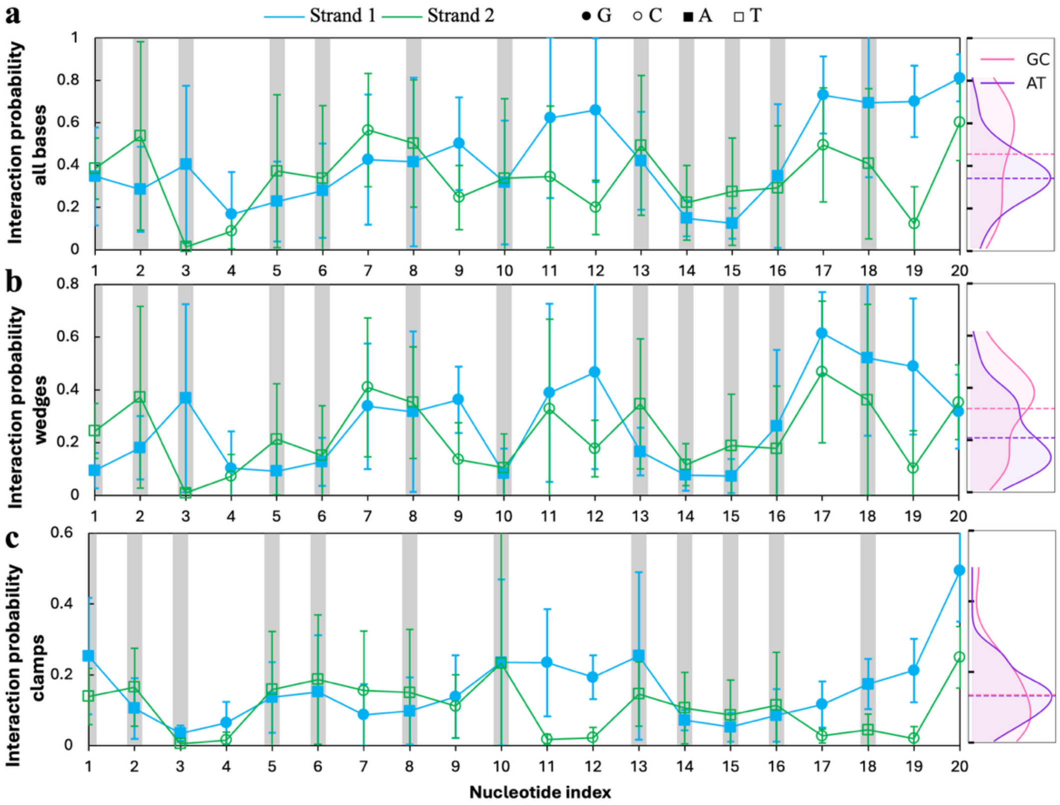


**Figure S9. DNA sequence-specific binding of protamine.** Same as Figure 3a-c, but for the 40%-GC 20-bp DNA.

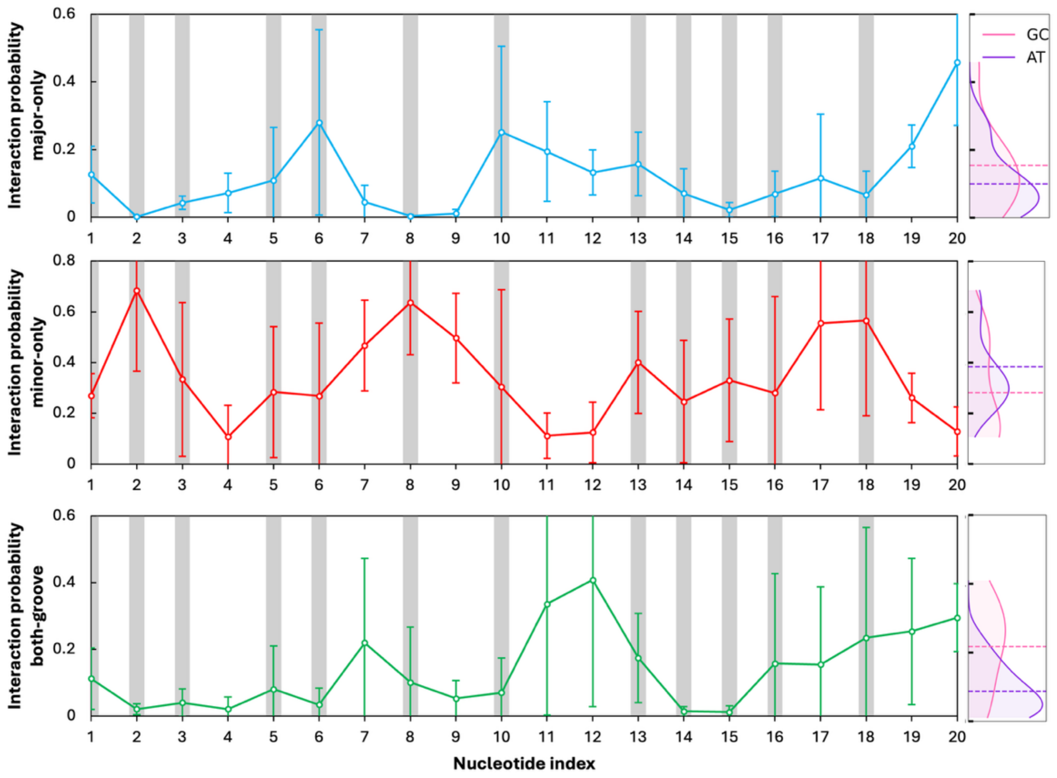


**Figure S10. Groove-specific Arg-interaction probabilities of bp along the DNA sequence.** Same as Figure 4c, but for the 40%-GC 20-bp DNA.

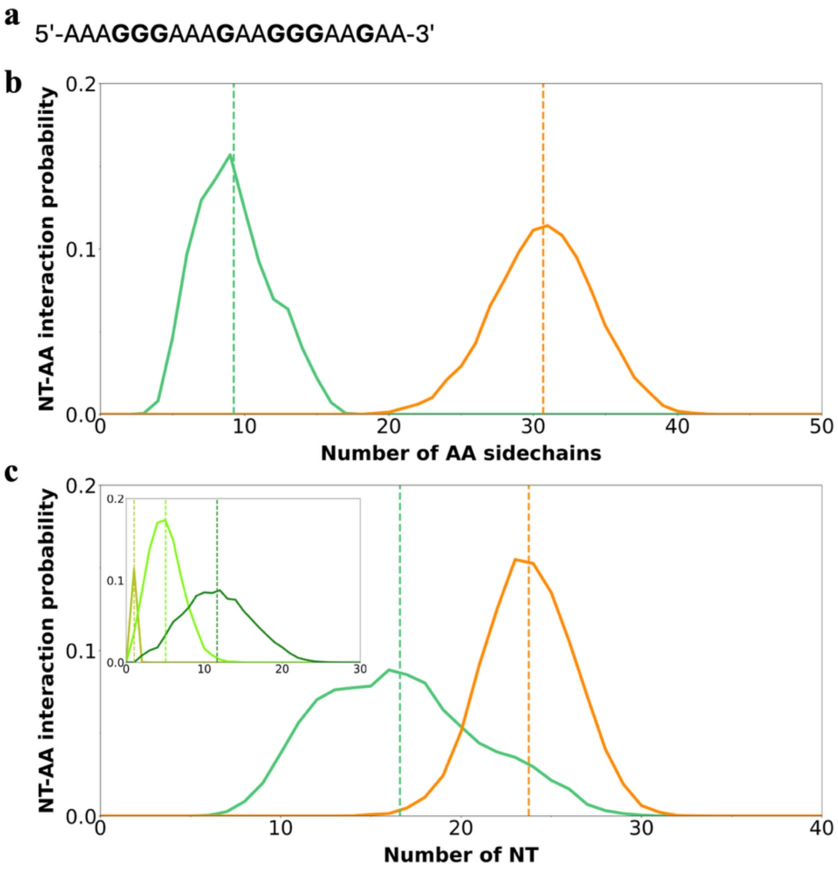


**Figure S11. DNA sequence-specific binding of protamine.** Same as Figure S8a-c, but for the 40%t-GC 20-bp DNA.

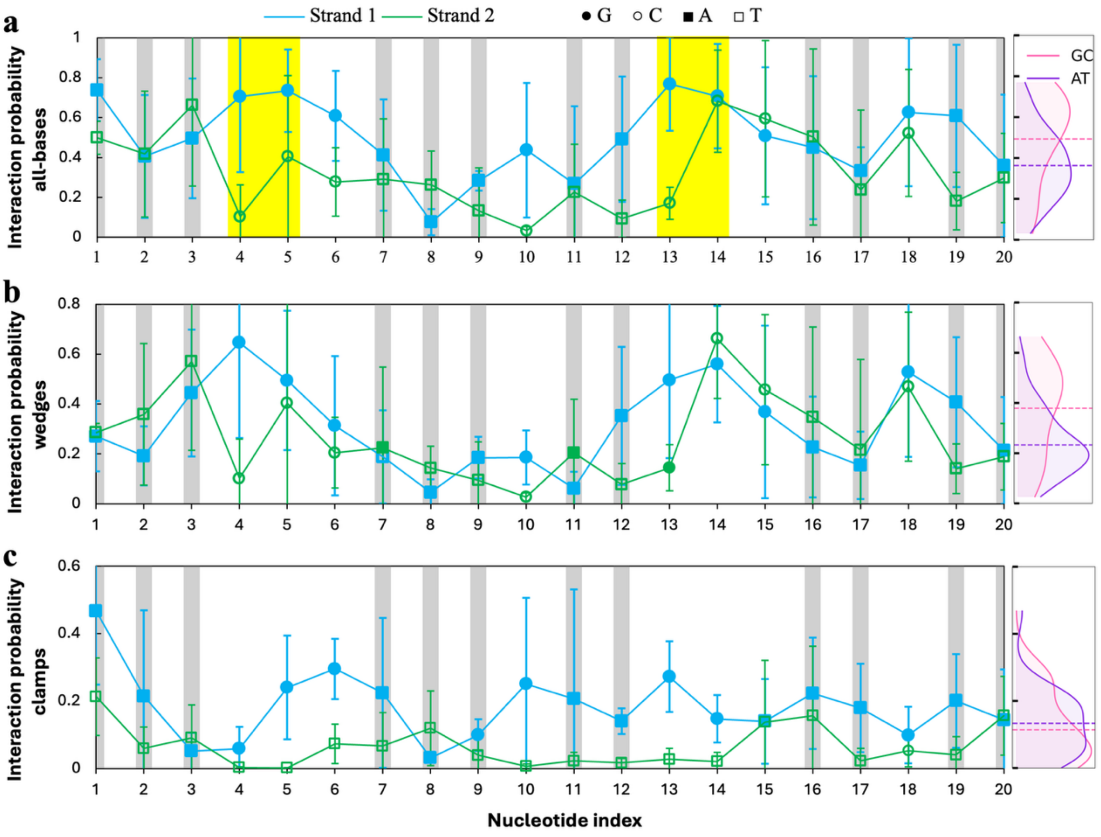


**Figure S12. DNA sequence-specific binding of protamine.** Same as Figure S9a-c, but for the 40%t-GC 20-bp DNA.

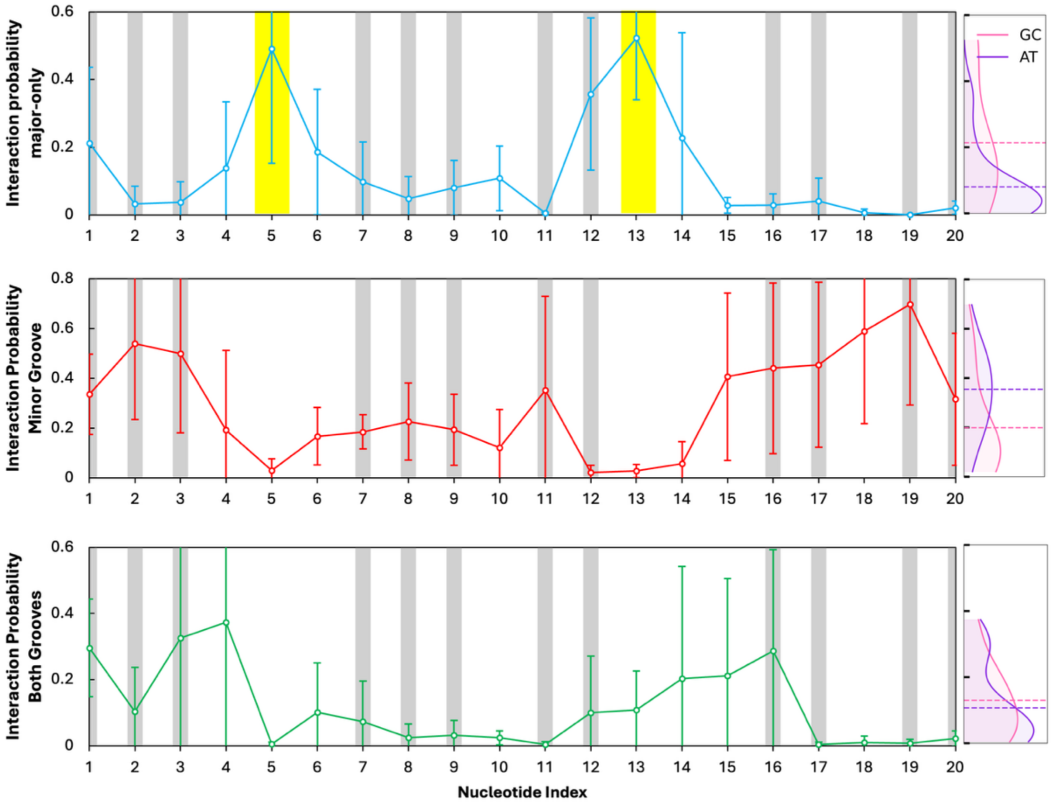


**Figure S13. Groove-specific Arg-interaction probabilities of bp along the DNA sequence.** Same as Figure S10, but for the 40%t-GC 20-bp DNA.

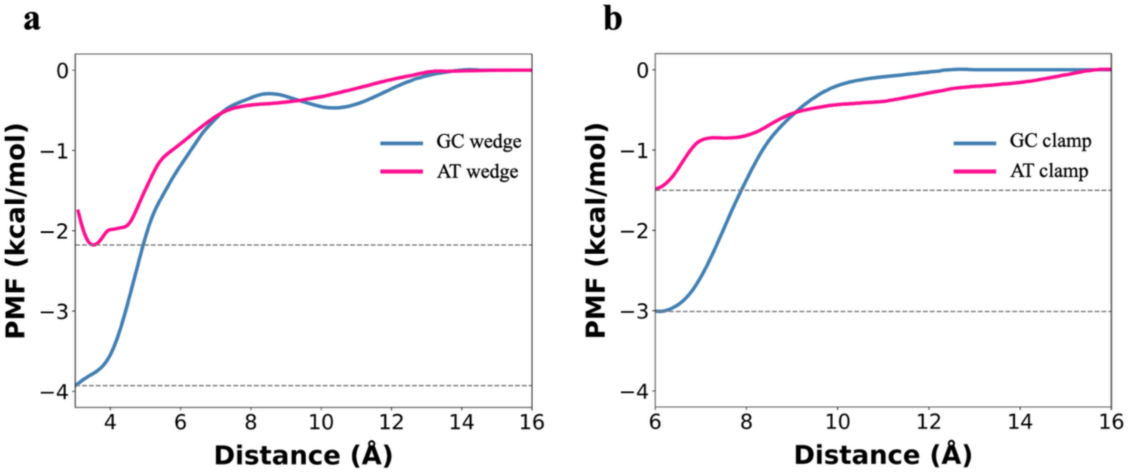


**Figure S14. Potentials of mean force for Arg-DNA interactions.** Same as Figure 5b, c, but with results for a wedge and a clamp, both involving two thymines at the major groove, added. KCl is at 0.15 M. Dashed horizontal lines indicate the free-energy minima in each case.

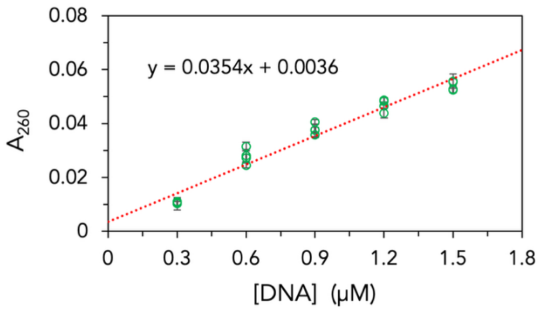


**Figure S15. The standard curve for converting absorbance at 260 nm to DNA concentration.** $A_{260}$ values are not distinguishable among the four 20-bp DNA sequences and used together to fit a linear standard curve. Error bars represent SDs among four replicate measurements for each DNA sequence at a given concentration.

**Movie S1. The traversal of an Arg side chain along a G-tract.** The G-tract comprises guanines at positions 22-24 on Strand 1 of the 35-bp DNA. Hydrogen bonding is indicated by dark dashed lines. The movie clip shows the 727-805 ns portion of one of the four 1000-ns replicate simulations.
